# PhycoMine: A Microalgae Data Warehouse

**DOI:** 10.1101/2021.09.27.462046

**Authors:** Rodrigo R. D. Goitia, Diego M. Riaño-Pachón, Alexandre Victor Fassio, Flavia V. Winck

## Abstract

PhycoMine is data warehouse system created to fostering the analysis of complex and integrated data from microalgae species in a single computational environment. The PhycoMine was developed on top of the InterMine software system, and it has implemented an extended database model, containing a series of tools that help the users in the analysis and mining of individual data and group data. The platform has widgets created to facilitate simultaneous data mining of different datasets. Among the widgets implemented in PhycoMine, there are options for mining chromosome distribution, gene expression variation via transcriptomics, proteomics sets, Gene Onthology enrichment, KEGG enrichment, publication enrichment, EggNOG, Transcription factors and transcriptional regulators enrichment and phenotypical data. These widgets were created to facilitate data visualization of the gene expression levels in different experimental setups, for which RNA-seq experimental data is available in data repositories. For comparative purposes, we have reanalyzed 200 RNA-seq datasets from *Chlamydomonas reinhardtii*, a model unicellular microalga, for optimizing the performance and accuracy of data comparisons. We have also implemented widgets for metabolic pathway analysis of selected genes and proteins and options for biological network analysis. The option for analysis of orthologue genes was also included. With this platform, the users can perform data mining for a list of genes or proteins of interest in an integrated way through accessing the data from different sources and visualizing them in graphics and by exporting the data into table formats. The PhycoMine platform is freely available and can be visited through the URL https://PhycoMine.iq.usp.br.

## Introduction

Microalgae have recently received a higher attention from the scientific community due to the several possible sustainable biotechnological applications of these organisms and their bioproducts. Since the release of the genomic sequence of the model species *Chlamydomonas reihardtii* (Merchant, et al., 2007), several other microalgae genomes have been sequenced and further functional genomics data have been generated mainly through omics approaches, contributing to significantly expanding our knowledge about these organisms and their evolutionary and metabolic characteristics.

Several databases have been recently developed to storage and mining microalgae molecular data, such as gene co-expression data (Aoki, et al., 2016), protein structural and physicochemical information (Kurotani, et al., 2017), enzymes annotation (Misra, et al., 2016), co-expression networks (Ferrari, et al., 2018), metabolic pathways (Schlapfer, et al., 2017; Zheng, et al., 2014), among others. However, with the increased availability of biological data, specially from the analysis of omics datasets, there is a current need for integrative databases that may contribute to a better data mining and visualization of microalgae information. Therefore, we have developed PhycoMine (https://PhycoMine.iq.usp.br), which is a database with biological information for microalgae integrating different data sources. PhycoMine was developed based on the InterMine platform, a biological database warehouse management tool, and we had implemented additional modifications and applications that permit a better analysis of the stored information. Biological data from different sources were processed to allow integration into the database, including data from the Phytozome repository for the annotated genes and transcripts, and Uniprot data for the information of annotated proteins.

In this first version of the PhycoMine platform, data integration is available for two microalgae species: *Chlamydomonas reinhardtii* and *Chlorella vulgaris*. The main types of data integrated are annotated genes, proteins, transcripts, gene families, gene ontology, gene orthologs and data from different RNA-seq and protein experiments, among other types of basic information, such as unpublished data for patterns of classical physiological responses such as cell growth, nitrogen and phosphate consumption profiles.

Several data classes were added in the database model of the InterMine platform to broadening and save new information types that are necessary for studying the two organisms mentioned and other to be included, and based in this amplified model, we developed several tools and graphs to analyze this new information individually and collectively.

The way to use the platform and the tools that can be found in it are briefly explained in this present work, however, we are going to concentrate in detailing the new classes of the model and the tools developed for data mining and visualization using this new information inside the database. For the use of the PhycoMine platform in its fullness, we advise the developers and end users to read the basic concepts of the platform InterMine (Kalderimis et al. 2014) through the documentation available at InterMine (http://intermine.org) webpage.

### Data in PhycoMine

Among all the data that is integrated in PhycoMine, we have relevant information about the annotation of genes, proteins, and transcripts, integrating all this knowledge with data of Gene Ontology, KEGG pathways, scientific publications, and the respective gene sequences, giving us a broad view of all the integrated information inside one element. In addition, the information of the gene expression levels in different experiments and their original data source and publication are connected. Same way, we added data from gene orthologs, gene families, and genomic position to be integrated with all the information stored in the platform. Proteomics data and basic cellular phenotypes such as growth and time-resolved nutrient consumption are also included for contributed datasets. The structure of the platform is gene centered and integrates several data types to a gene or list of genes of interest that are the initial query point.

### Data Sources

The information stored inside the database comes from different sources and external data repositories and were pre-processed before storage. InterMine has developed the structure on how to introduce the data of genes, transcripts, and proteins with their sequences, using GFF3 files, the FASTA file of the genome of each organism, and the xml file for proteins. In PhycoMine, the GFF3 and FASTA files for *Chlamydomonas reinhardtii* were downloaded from the Phytozome database (version 12), and for the organism *Chlorella vulgaris*, the genomics data was retrieved in the database of *Chlorella vulgaris* Genome Project in the “Genome Project Solutions Database” (http://chlorella.genomeprojectsolutions-databases.com/). For both organisms, the xml file for proteins was obtained from UniProt (The UniProt, 2017).

To integrate the protein annotation obtained from UniProt with all the information obtained from both Phytozome and “Genome Project Solutions Database”, a workflow was developed in which the sequences of the proteins are compared with the proteins of the other two databases using BLAST (Altschul, et al., 1990). The BLAST result was merged with the xml file, in order to generate a new xml file with the necessary data to integrating with the genome data.

We developed new scripts and/or workflows for data formatting and integration of the data annotation for genes, transcripts, and proteins with the information from Gene Ontology (Ashburner, et al., 2000), pathways, gene families, orthologs and expression levels data as well as scientific publications. For Gene Ontology, it was necessary to create a new table format file that integrates the genes and the ontology file (.obo) within the PhycoMine. The pathways data were downloaded from the KEGG database, processed into two files, one with the information of all the pathways, and the other with the relationship between the genes. Gene Families information was obtained from the eggNOG (Huerta-Cepas, et al., 2019) and PlnTFDB (Perez-Rodriguez, et al., 2010) databases and linked to a gene and/or gene list. The gene orthologs were annotated for the protein sequences FASTA file using the orthoFinder software (Emms and Kelly, 2019).

Gene expression data was retrieved for several different experimental conditions from the NCBI’s SRA database (see Supplementary data 1 for the list of accessions) and a workflow was developed in such a way that a matrix with the expression values of each gene in the different experiments were included in the platform.

With the downloaded SRA files, the fastq-dump option of sra toolkit was executed, then the information was processed within BBduk2 (https://jgi.doe.gov/data-and-tools/bbtools/) for data pre-processing, resulting in a fastq.gz file, which with it, could be processed inside of the Salmon software giving us the results of the expression levels for each experimental condition available. Finally, a script developed in house in R language, processed the data by tximport function, generating the resulting file in a table format that contains the levels of genetic expression and can be included within the PhycoMine.

### Architecture

To store the novel information, it was necessary to expand the InterMine model, with new classes been created to describe the elements that correspond to the new information. Classes were added for the type of experimental data (e.g., transcriptomics or proteomics data), description of the experiments and their conditions and the gene expression data obtained from the NCBI database. Gene family, KEGG pathways, orthologs and gene expression networks (ranges and clusters) were included in the platform as new data classes and categories as shown in Fig. 1

**Fig. 1.**
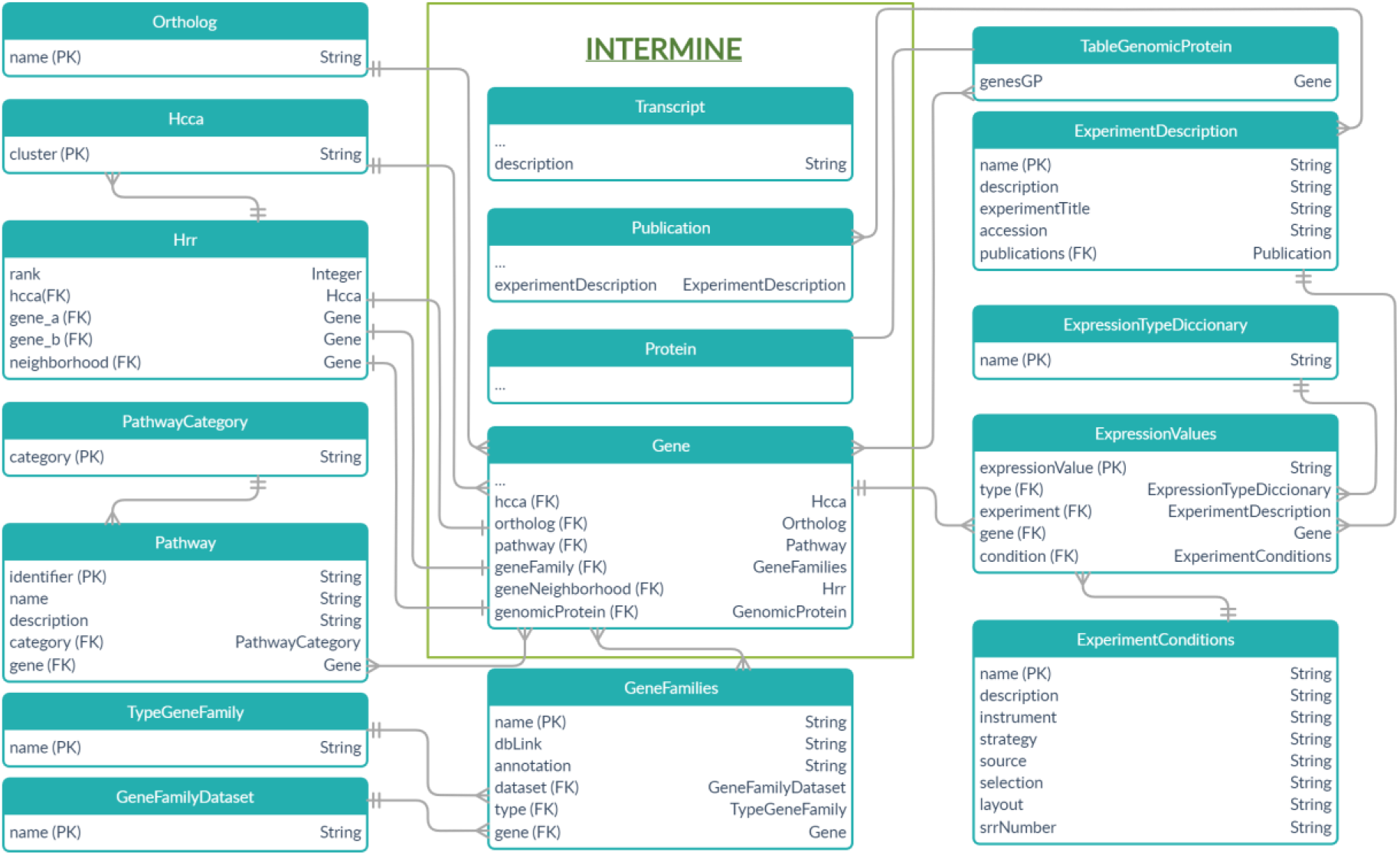
Structure of the extended database model of PhycoMine. The featured warehouse structure of the PhycoMine database is an extended version of the InterMine warehouse structure. New data classes were included and connected considering a gene centered hierarchy of linkages.

### How to Search Information

There are several ways to access the information stored in the database, either to obtain data from a single element or from a list of them. From the home tab, a specific search tool is available for input gene, protein, and keywords, that allow the users to perform a direct search using gene or protein identifiers, or several of them separated by commas.

Importantly, the identifiers accepted into the PhycoMine are originals from Phytozome model added of the version of the genome annotation adopted at the current version of PhycoMine database (v5.5). Therefore, the users should add the suffix **“**.**v5**.**5”** to any gene or protein ID used as input data for search purposes; for instance, Cre17.g734450.**v5**.**5**, where the .v5.5 is the suffix.

The search retrieves a list of elements found in the database that match to the input identifiers. Likewise, the users can build custom queries in the QueryBuilder tab, selecting the classes of interest to start the query and then built it with joints and constraints features. There are also predefined query templates for executing common queries faster. Finally, the user can search for genomic elements giving the genome region of interest. All these search functionalities can also be executed programmatically from another external application using an Application Programming Interface (API).

Once the search is executed, the results show the details of the queried element(s) in the object page. Further analysis of the group of elements of interest can be performed by adding the results to a list. The lists can only be of one type of object (e.g., genes), and these will be saved in the new created list.

To save the queries that were executed, the created lists and usage history, it is necessary to register and logged in into the system with a new account in MyMine tab.

### Data mining and visualization

To exemplify the possibilities of data mining, display and system visualizations, we created a list of identifiers that included the differentially expressed genes from the work of Gallaher et. al., 2018, that was investigating nitrogen deprivation data (Gallaher, et al., 2018). The list of differentially expressed genes summed 734 entries. We extracted a subset of 50 genes out of the 734 for creating an example input list of genes for demonstrating the types of data visualization and analysis available in PhycoMine.

### Lists

Data mining can be performed by using lists. A list of objects (genes or proteins) can be inserted into the Analyze search option at Home page or obtained from the PhycoMine database by accomplishing certain constraints. In the case of a gene list, a series of visualizations are generated by default that contributes to mining several different information about the group of objects in that list. In one of the visualization widgets a heatmap is displayed showing the gene expression behavior of the input genes during a given experiment or comparison of experiments (Fig. 2A) and, next to it, the gene profile analysis is shown, which also compares the expression level of all genes in the input dataset, showing the mean gene expression profile for each gene in each experimental condition compared (Fig. 2B). Gene expression results from two different experiments can be directly compared using the visualization tools.

**Fig. 2.**
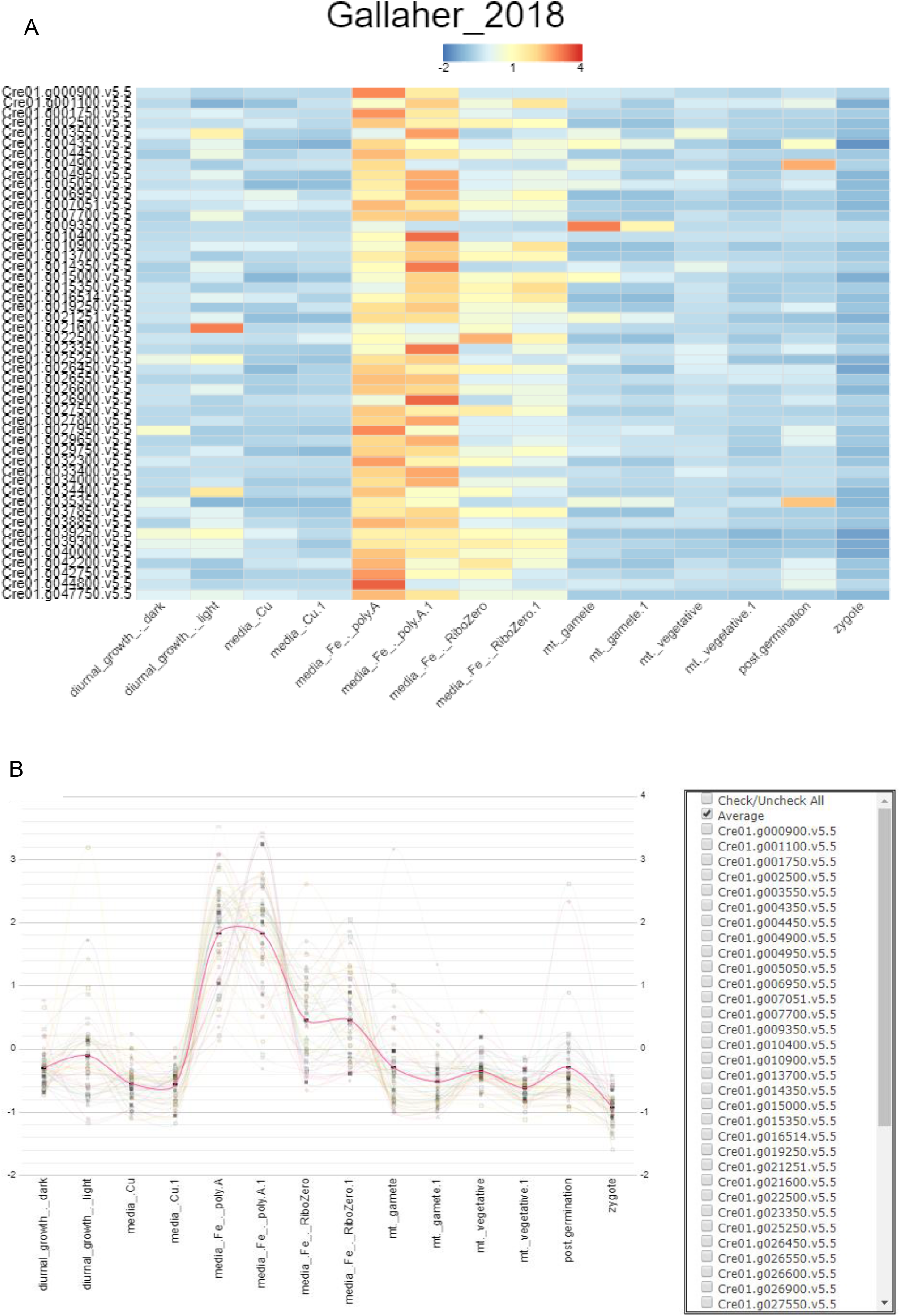
Gene expression visulization. (A) Heat map of the gene expression levels of the selected 50 genes extracted from Gallaher experiment. The gene expression data was retrieved from SRA archive and data analysis was performed for all datasets using a common pipeline for data pre-processing and differential expression analysis. Gene expression values were mean centered, and a color scale bar is representing the calculated z-scored abundance of each gene for the different experiments; (B) Visualization of the gene expression level of selected genes in different experiments. The mean gene expression level of the 50 selected genes is shown as dispersed circles for the different experimental conditions indicated in the y axis. The expression profile of the individual genes of interest can be selected and filtered in or out of the multiple comparison. A trend line is indicated for visualization purposes and the bold red curve is indicating the mean value of the gene expression for the 50 selected genes.

In our example input dataset (50 genes), we can see the pattern of gene expression variation during the transition between the experimental conditions of the study performed by Gallaher et al. 2018.

The second widget and visualization tool included by default in PhycoMine is a scatterplot, which shows the comparison of the expression levels of each gene in two different experimental conditions, from two different studies. In this case, the comparison was made between different conditions that were described in two experiments (independent studies), with each node equivalent to each gene in the list.

In our example, we compared two datasets: the expression data of the 50 selected genes from the study performed by Gallaher et. al. 2018 with the data generated by the experiment performed by Voshall et. al. 2015 (Voshall, et al., 2015) displaying the data in a Scatter plot, where more distant elements are the ones that contain a higher variability between the two studies. **Fig. 3**

**Fig. 3.**
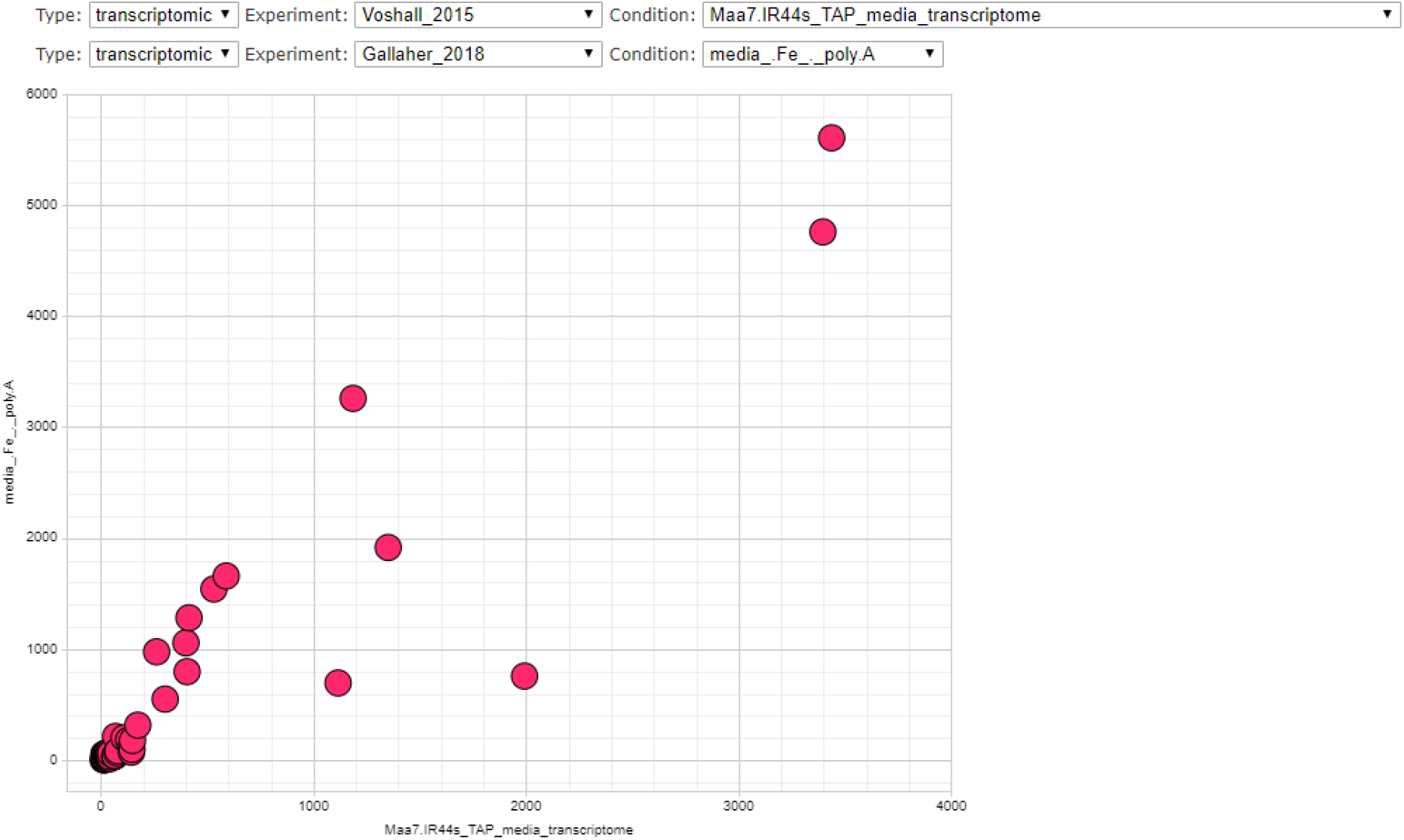
Scatterplot of the gene expression level of selected genes from two independent transcriptome studies. Gene expression data from the two different experiments were reanalyzed using the same transcriptome data analysis pipeline and the visualization of the comparison of the experiment performed by Gallaher and colleagues and the experiment done by Voshall and colleagues. The graphical visualization of the two experiments is representing the gene expression values of the 50 genes selected for demonstration purposes.

Additionally, we have included widgets for functional annotation enrichment analysis, which the results are the number of times an object appears in the list and the p-value that is the probability that a result occurs by chance, thus a lower p-value indicates greater enrichment. This analysis can also be performed for enrichment analysis based on publications, gene ontology (Ashburner, et al., 2000), KEEG (Kanehisa and Goto, 2000) and gene families divided between those belonging to EGGNog (Huerta-Cepas, et al., 2019) and PlnTFDB (Perez-Rodriguez, et al., 2010). For metabolic data obtained from KEGG, two enrichment analyzes were developed: one containing pathways of diseases that do not belong to the microalgae and other that does not have these routes (**Fig. 4**).

**Fig.4.**
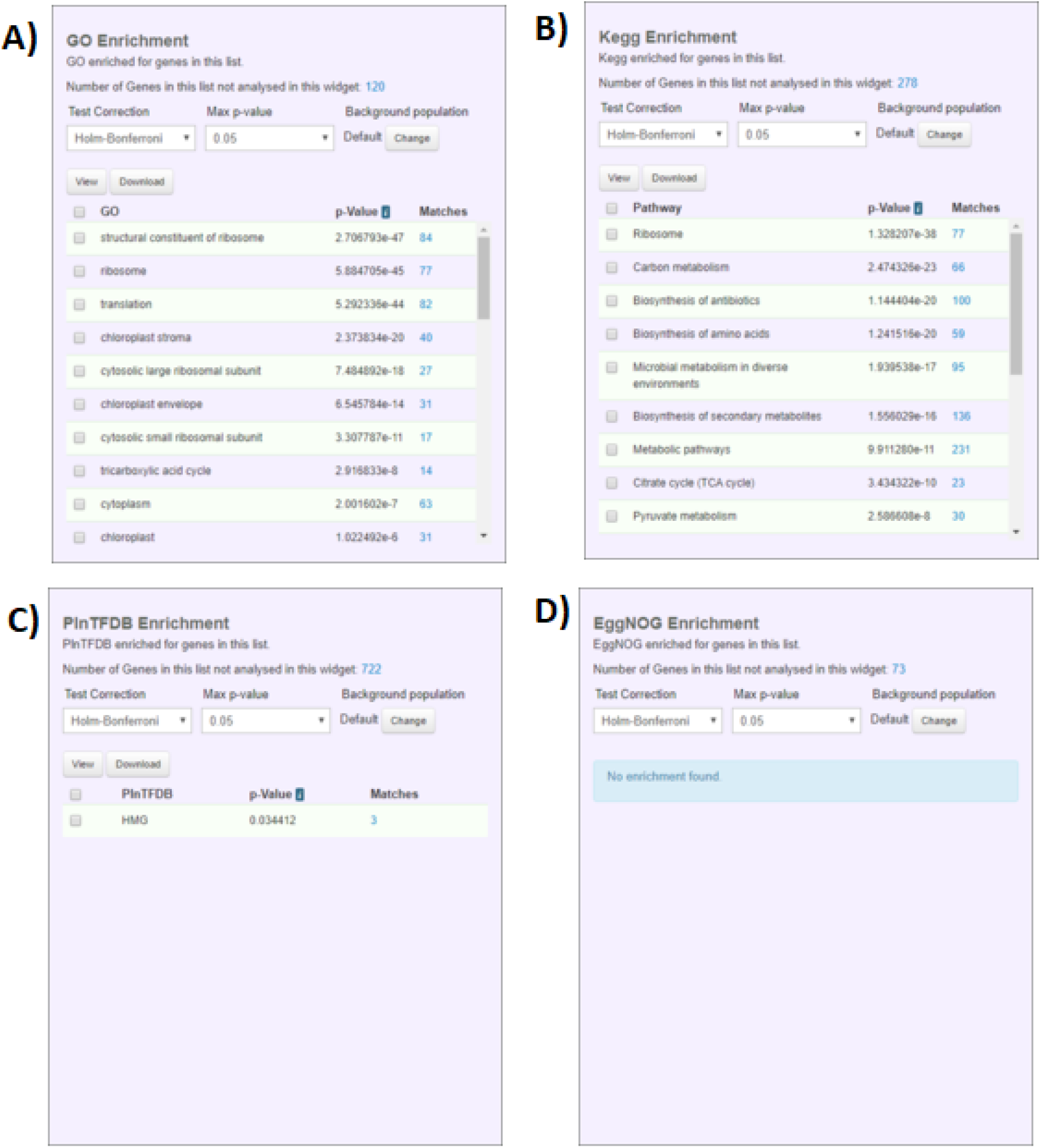
Functional annotation widgets present in the PhycoMine. Enrichment graphics show which objects appear with statistical significance with respect to a set of genes that represent the total universe or background. The results of the searches for the example input list using the widgets are shown in separated tabs, as follow: A) Gene Ontology enrichment; B) KEGG enrichment; C) Transcription factors (PlnTFDB) enrichment and D) EggNOG Enrichment.

### Object

The object page for an element contains all the annotation information of the matched object, related data, and the integration with other internal or external databases is retrieved and it has additional classes. In the elements of the gene type, one can see various visualization tools that contribute to a better analysis of the gene function. Among those visualizations we have the Gene Atlas, a visualization in the form of a bar graph that shows the expression level of that selected gene in an experiment. Within that visualization, you can choose the scale and order of the conditions, as well as the experiment which you want to visualize. The comparison between two experiments is included in this tool, which shows the two names of the conditions compared and a scatterplot graph showing a different view in the comparison of the two conditions.

For exemplification purposes, we decided to show the tools of the genes object page for the Cre01.g021600 gene from *Chlamydomonas reinhardtii*. In the Fig. 5A we can see the level of expression of the gene Cre01.g021600 in the study performed by Gallaher and colleagues (Gallaher, et al., 2018) (Fig. 5A), then the same standardized data (applied Z-score) is observed in Fig. 5B. In the third part, three conditions of the Gallaher experiment are compared with three other conditions of the Voshall experiment (Voshall, et al., 2015) (Fig. 5C), and finally, the scatter plot display option is shown, where it compared equally way three conditions of the Gallaher experiment with the Voshall experiment (Fig.5D).

**Fig.5.**
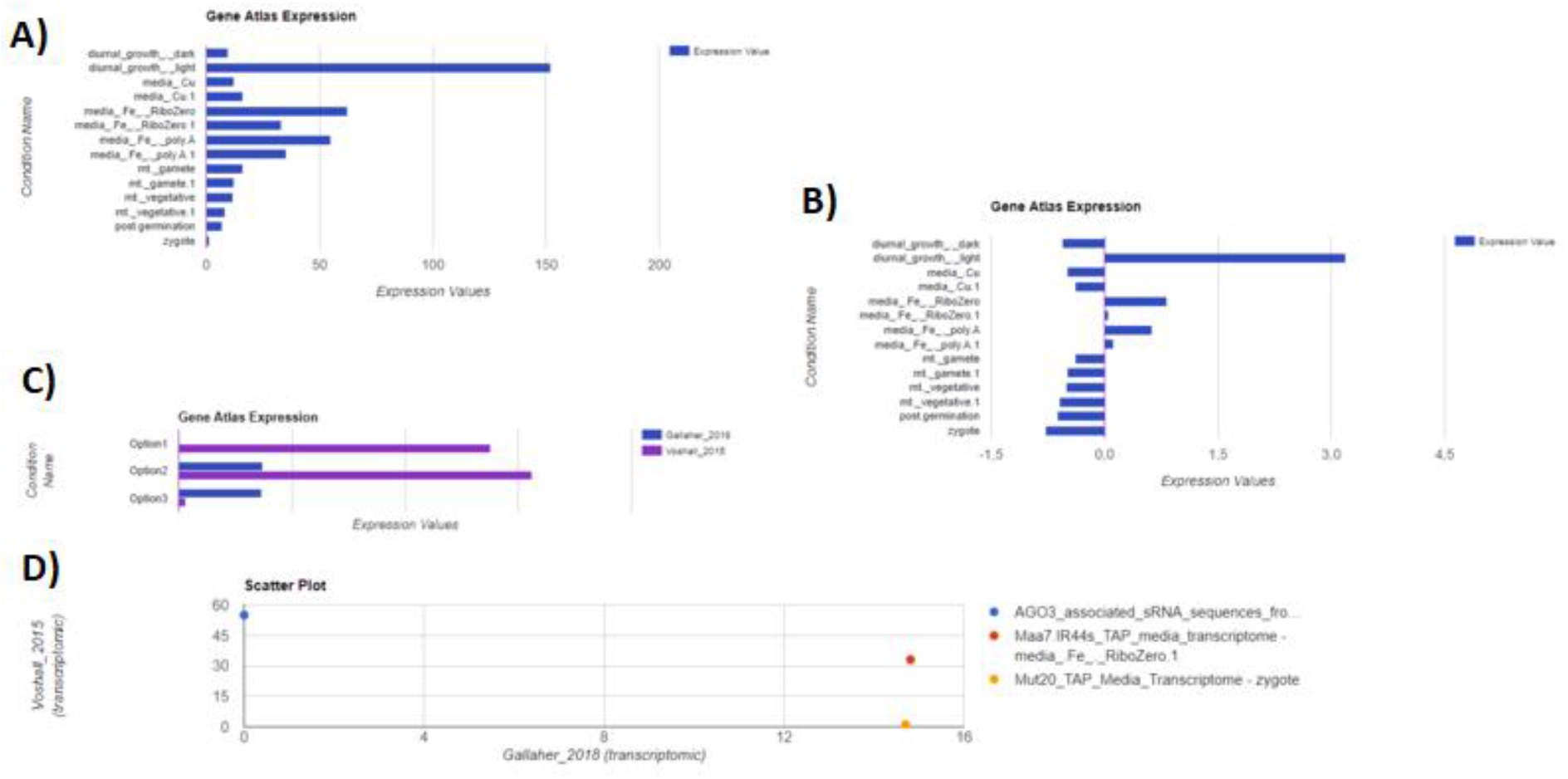
Gene centered expression information and comparative analysis. The level of gene expression reported in the selected conditions of the experiments performed by Gallaher are shown as bar graphs. A) expression levels for gene Cre01.g021600 order by the experimental conditions selected; B) z-score calculated expression levels for gene Cre01.g021600 order by the experimental conditions selected; C) expression levels for gene Cre01.g021600 in Vorshall and Gallaher experiments; D) scatterplot comparing the expression levels for gene Cre01.g021600 under conditions of the experiments Gallaher and Voshall.

Another data analysis tool that is available on the object page is the Co-expression Networks analysis, a tool that has two gene networks generated with a heuristic clustering algorithm (Mutwil, et al., 2010). The algorithm returns two important matrices to generate networks, neighborhood and clusters. With both, you can calculate the two networks generated in the application. The tool lets us highlight within the network, the orthologs of each gene and visualize them by gene families; it can also represent the size of the node of each gene to represent the expression of that gene in a selected condition.

In the Co-expression Networks list (Fig.6 A), it is possible to either create a list of genes that belong to the co-expression network or open the network visualization tool. In the first network of co-expression (Fig.6 B), the neighborhood of the gene is showing all the genes that are co-expressed with the gene that was calculated, in this case, all the nodes having a connectivity with the node that represents the Cre01.g021600 gene. In this case, the user has some options to customize the network features (e.g., colors, node formats, etc.), where the color of the nodes represents the orthologs associated to the queried gene, the node form differentiates the gene families, and the size is directly proportional to the expression of that gene in a selected experimental condition. Finally, we have in the last image the co-expression network of the cluster to which the selected gene belongs (Fig. 6C). It is not necessary that all genes have a connectivity with a specific node, but they are all genes that have a similar behavior between the experiments, as can be seen, all the display options were turned off, only the level of connectivity between the nodes is shown. The more attached to the green color, the better connection between the nodes exists, on the other hand, if the connection is weak, the node is red. Connection levels are measured by ranges, from 0 to 49, therefore, the lower the range, the more correlated the nodes are (Fig.6).

**Fig. 6.**
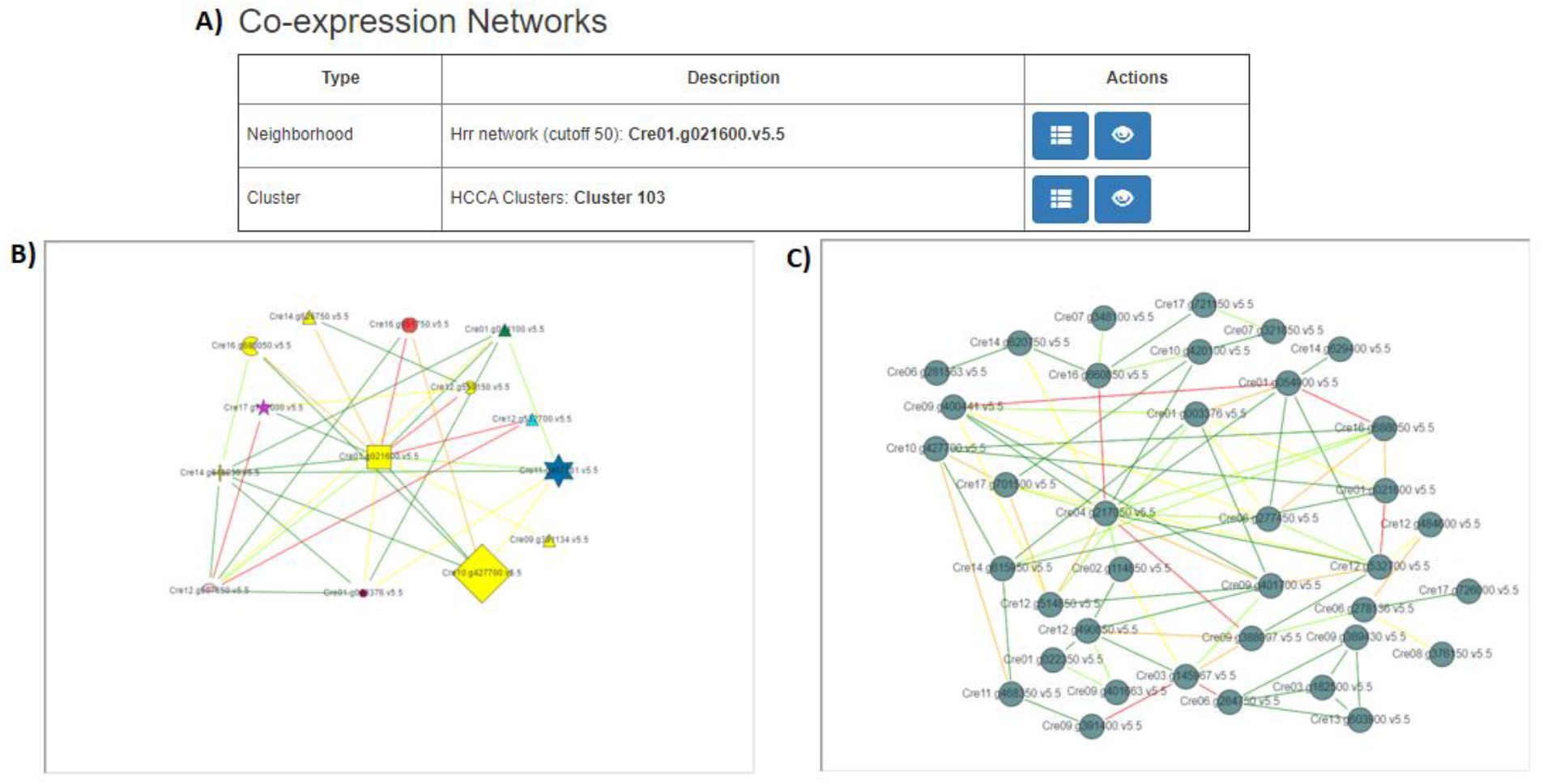
Online Co-expression Network Analysis in PhycoMine. A) Co-expression network types, with the options to create the list with the genes to be presented in the network, and the option to visualize the network. B) Neighborhood network graph; C) Cluster Network of the gene Cre01.g021600.v5.5.

For tracking the genes by their genomic coordinates and position at the genome level, the JBrowse tool was also integrated into PhycoMine to visualize the gene in the genome with all its integrations (Buels, et al., 2016).

As mentioned earlier, the gene family information was added integrated to the information of gene object, allowing to retrieve the data source eggNOG (Huerta-Cepas, et al., 2019) or PlnTFDB (Perez-Rodriguez, et al., 2010)) and the Scope dataset to which it belongs, together with an annotation and a link that leads to the gene family page for both eggNOG or PlnTFDB, generating a list of gene families related to the selected gene Cre01.g021600 (Fig. 7).

**Fig. 7.**
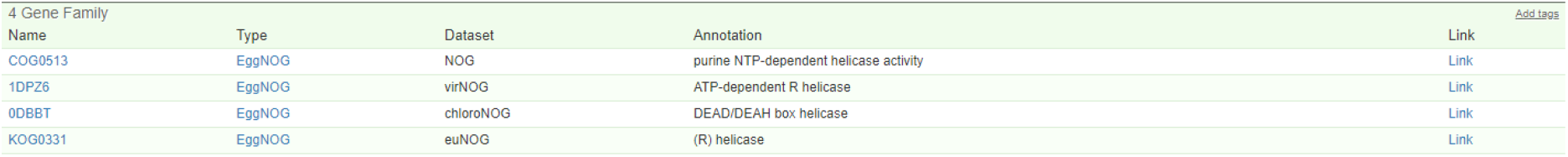
List of gene families of the gene Cre01.g021600. Data type, annotation and the link of data source can be listed for a selected object.

In a similar way, the genes that belong to the same group of orthologs of the gene being studied are shown, and on the same page, the metabolic pathways to which it belongs. (Fig. 8)

**Fig. 8.**
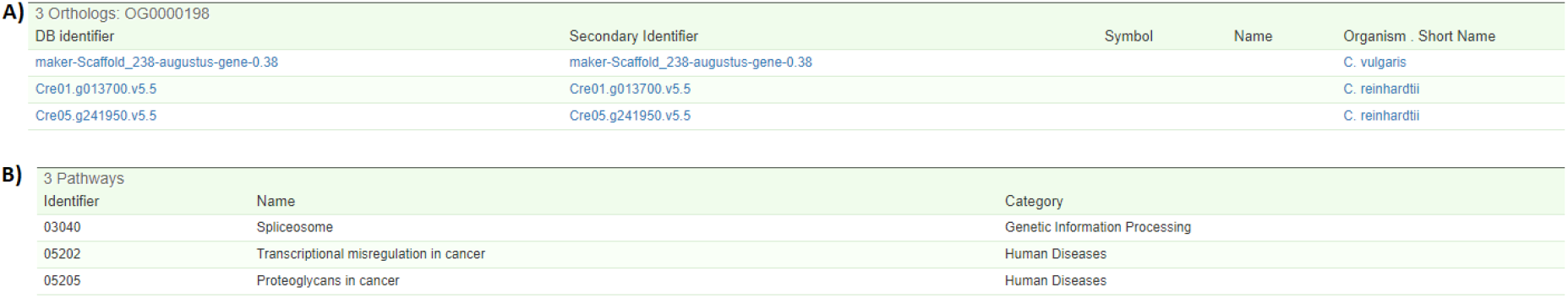
Orthologous genes data integration. A) List of the genes that belong to the same ortholog group as the Cre01.g021600 gene; B) List of metabolic pathways to which the Cre01.g021600 gene belongs.

## Discussion

The PhycoMine platform was built based on InterMine (http://intermine.org) structure which contributed to the organization and integration of biological data from the microalgae *Chlamydomonas reinhardtii* and *Chlorella vulgaris*. In addition, we developed new forms and scripts for the insertion of new information or new relationships between the data within the database, in order to have a broader and integrated vision of the biological information.

In the same way, the introduction of RNA-seq experimental data to the platform was implemented. In this case, a workflow was developed to analyze the information coming from NCBI (https://www.ncbi.nlm.nih.gov) using the same data analysis pipeline and parameters and, in addition, having greater efficiency at the time spent on the analysis. The development of the scripts for analysis and alignment of the data was very important when inserting information into the database, for RNA-seq, orthologs and gene families.

Finally, with this processed information, graphs are shown to the users of the system at different levels or classifications. Placing the necessary tools in a structure that is facilitating the data search by non-specialized professionals.

All the PhycoMine code, as well as the analysis and alignment scripts, and the data of the microalgae *Chlamydomonas reinhardtii* and *Chlorella vulgaris* can be found at Github repository labisapplication (https://github.com/labisapplication/Phycomine-features-Labis).

